# Lead as a toxic environmental toxicant in models of synucleinopathies

**DOI:** 10.1101/2024.10.10.617554

**Authors:** Liana Shvachiy, Ângela Amaro-Leal, Filipa Machado, Isabel Rocha, Vera Geraldes, Tiago F. Outeiro

## Abstract

Lead, a toxic heavy metal, is prevalent in various industrial applications, contributing to environmental contamination and significant health concerns. Lead affects various body systems, especially the brain, causing long-lasting cognitive and behavioral changes. While most studies have focused on continuous lead exposure, intermittent exposure, such as that caused by migration or relocations, has received less attention. Importantly, lead exposure intensifies the severity of Parkinson’s disease (PD) and dementia with Lewy bodies, diseases involving the accumulation of alpha-synuclein (aSyn) in the brain and in the gut. Although, the precise mechanisms underlying these observations remain unclear, oxidative stress and mitochondrial dysfunction likely play a role. Here, we investigated how two different profiles of lead exposure - continuous and intermittent - affect models of synucleinopathies. We found that lead exposure enhances the formation of aSyn inclusions, resulting in an increase in both their number and size in cell models. In addition, we found that animals injected with aSyn pre-formed fibrils display serine 129-phosphorylated aSyn inclusions and a reduction in astrocytes in the substantia nigra. These animals also display neuronal damage and alterations in locomotor activity, exploratory behavior, anxiety, memory impairments and hypertension. Our results suggest a mechanistic link between environmental lead exposure and the onset and progression of diseases associated with aSyn pathology. Understanding the molecular and cellular interactions between lead and aSyn is crucial for shaping public health policies and may provide novel insight into strategies for mitigating the impact of environmental toxins on neurodegenerative processes involved in Parkinson’s disease and related synucleinopathies.

**Key points summary:**

- Lead exposure increases the number and size of alpha-synuclein (aSyn) inclusions in cell models.∼
- Animals exposed to aSyn fibrils and lead show increased phosphorylation of aSyn, neuronal damage and reduced astrogliosis in the *substantia nigra*.
- Lead-exposed animals exhibit impaired locomotion, anxiety, memory deficits, hypertension and chemoreceptor reflex hypersensitivity.
- The study suggests a mechanistic link between environmental lead exposure and the progression of diseases involving aSyn pathology, such as Parkinson’s disease.

## Introduction

Lead, a naturally occurring heavy metal (chemical element Pb), is used in a variety of widely used items such as automobiles, paints, and traditional cosmetics. Its broad usage has resulted in environmental damage, particularly contamination of drinking water, resulting in human exposure and, thereby, in public health consequences (World Health Organization, 2010, 2021; Wani *et al*., 2015; Sani & Amanabo, 2021; Tempowski & World Health Organization, 2021). Lead toxicity is a major environmental health concern, with the World Health Organization estimating a loss of around 1 million lives, in 2019, due to lead exposure (World Health Organization, 2021). There are two forms of lead exposure: (i) occupational or acute exposure and (ii) environmental or chronic exposure, with the latter being more dangerous in the long term (World Health Organization, 2010, 2021; Organization, 2022). Surprisingly, new environmental lead exposure profiles have been recently described, including the intermittent lead exposure profile that has been extensively studied and characterised in our laboratory in recent years, which has different health effects to those that have been extensively studied over several years (chronic/permanent lead exposure) (Shvachiy *et al*., 2018, 2020a, 2022, 2023).

Lead is primarily found in soft tissues such as the brain, liver, and kidneys and accumulates in bones and teeth for long-term storage. In women, lead is released during pregnancy, exposing the foetus (Sani & Amanabo, 2021). Children are particularly vulnerable to lead toxicity as they absorb 4-5 times lead than adults (Laura Mayans, n.d.; Academy *et al*., 2005). The most serious consequences are a reduction in cognitive performance and behavioural problems in children (Koller *et al*., 2004; Academy *et al*., 2005). Adults may experience haematological consequences such as anaemia, gastrointestinal disturbances such as stomach colic, and renal dysfunction (Todd *et al*., 1996; Flora *et al*., 2012; D. A. Gidlow, 2015; Wani *et al*., 2015). However, the major health impacts include cardiovascular dysfunction, an elevated risk of hypertension, ischemic heart disease, and stroke, along with neurological disturbances like headaches, irritability, depression, lethargy, convulsions, muscle weakness, ataxia, tremors, and hearing loss (Vaziri, 2008; Virgolini & Aschner, 2021). Other major health consequences of lead include neurotoxicity and, more particularly, neurodegeneration, as it has previously been linked to numerous neurodegenerative disorders such as Alzheimer’s disease (AD), Amyotrophic lateral sclerosis, and Parkinson’s disease (PD) (Toscano & Guilarte, 2005; Wu *et al*., 2008; Mason *et al*., 2014; Waseem *et al*., 2014; Chin-Chan *et al*., 2015; Bihaqi *et al*., 2018; Rocha & Trujillo, 2019).

PD is a complex, chronic, and progressive neurodegenerative illness characterized by typical motor symptoms such as resting tremor, stiffness, postural instability, and bradykinesia (Spillantini & Goedert, 2000; Ogun, 2002; Radhakrishnan & Goyal, 2018). However, non-motor symptoms such as hyposmia, postprandial and orthostatic hypotension, supine hypertension, REM sleep behavior disorder, constipation, baroreflex dysfunction, depression, and dementia are also prevalent in PD (Spillantini & Goedert, 2000; Ogun, 2002; Fleming, 2011; Goldstein, 2013; Radhakrishnan & Goyal, 2018; Scorza *et al*., 2018; Chen *et al*., 2020; Elfil *et al*., 2020). Several of these non-motor symptoms arise early in the illness. However, the patient is diagnosed with PD only after the motor symptoms are evident (Braak *et al*., 2003; Hawkes *et al*., 2010)

PD belongs to a group of neurological disorders known as synucleinopathies that are characterized by the accumulation of intracellular protein inclusions rich in protein the alpha-synuclein (aSyn) (Murray *et al*., 2001; Goedert *et al*., 2017; Henderson *et al*., 2019a; Coon & Singer, 2020). Although the precise biochemical function of aSyn is unclear, it is thought to regulate the pool and release of synaptic vesicles in neurons (Iwatsubo, 2007; Marques & Outeiro, 2012; Wong & Krainc, 2017; Mehra *et al*., 2019). Nevertheless, aSyn is not only a neuronal protein and is also present in other tissues such as the heart, muscles, and blood (Marques & Outeiro, 2012; Pinho *et al*., 2019a; He *et al*., 2021).

A link between lead exposure and PD has already been shown, most notably in studies that found that occupational exposure to lead increase the risk of PD (Gorell *et al*., 1997; Coon *et al*., 2006; Weisskopf *et al*., 2010a; Parsian *et al*., 2021; Paul *et al*., 2021). Relatively few studies have focused on the impact of lead on aSyn (Zhu *et al*., 2011; Bihaqi *et al*., 2018). However, to our knowledge, the effect of lead on aSyn aggregation in cell and animal models, and associated phenotypic alterations, have not been studied. Here, we conducted a detailed characterization of the effects of two different lead exposure profiles in models of aSyn pathology. Ultimately, our goal was to obtain insight into molecular and functional effects that can form the basis not only for informed preventive public health measures, but also for possible strategies for therapeutic intervention.

## Methods

### In vitro studies

#### Preparation of recombinant human aSyn protein species

The preparation of recombinant human aSyn protein was performed using an established protocol (Al-Azzani *et al*., 2022). Briefly, the plasmid pET21-aSyn encoding for human aSyn was transformed into competent *E. coli* BL21-DE3 (Sigma, Germany), and cells were grown in 2x LB medium (Sigma, Germany) with 200g/mL ampicillin (Carl Roth, Germany) at 37°C with shaking. The expression was induced for 2 hours with 1 mM isopropyl-thiogalactopyranoside (IPTG, Apollo, UK) at OD600 0.5-0.6.

Before boiling for 15 minutes at 95°C, lysates were sonicated on ice for 5 minutes (30s on, 30s off pulses, 60% power). The supernatant containing aSyn was obtained after boiling the protein samples. Supernatants were purified using anion exchange chromatography (HiTrap Q HP, GE Healthcare) on Äkta Pure 25M (Cytiva) with a mobile phase of 25 mM Tris pH 7.6 and a linear gradient of 9 column volumes of elution buffer to 1M NaCl. Protein absorbance at 280 nm was measured, and aSyn concentration was calculated using the molar extinction coefficient 5960 M-1cm-1. To evaluate protein purity, 5 g of each sample was run on a 15% SDS-PAGE gel and stained with Coomassie dye.

To generate seed fibrils (PFF), 200 µM of monomeric aSyn solution was incubated in a 96-well plate at 37°C with 500 rpm shaking in the presence of Teflon polybeads. The fibrils were precipitated by ultracentrifugation for 1.5 hours at 15°C in an Optima MAX-TL (Beckman Coulter, Fullerton, CA) with a TLA-120.2 rotor at 80,000 rpm. The supernatant was carefully removed after centrifugation, and the pellet was washed twice with PBS. The resulting pellet of amyloid fibrils was resuspended in PBS after 1 minute of sonication in a bath-type ultrasonicator (Branson 1510, Danbury, CT). Before usage, the amyloid fibrils were kept at 4°C and re-sonicated for 1 minute.

### Cell cultures and treatments

#### Primary neuronal cultures

C57BL6/J#00245 wild-type E15.5 mouse embryos, obtained from the University Medical Center Göttingen animal facility (Göttingen, Germany), were used to prepare primary whole brain neuronal cultures, as previously described (Caldeira Brás *et al*., n.d.). Pregnant animals were sedated with carbon dioxide poisoning and the embryos were removed from the uterus. Following meningeal excision, the whole brain was dissected under a stereomicroscope and transferred to ice-cold 1x Hanks’ balanced salt solution (CaCl_2_ and MgCl_2_ free) (HBSS; Gibco Invitrogen, CA, USA) supplemented with 0.5% sodium bicarbonate solution (Sigma-Aldrich, MO, USA). The process was halted by adding 100uL fetal bovine serum (FBS; Anprotec, Bruckberg, Germany) and 100μL DNase I (0.5 mg/mL; Roche, Basel, Switzerland) after 15 minutes (min) of trypsinization (100µL of 0.25% trypsin; Gibco Invitrogen, CA, USA). After dissociation, the cell suspension was centrifuged at 300xg for 5 minutes before being resuspended in a pre-warmed neurobasal medium supplemented with 2% B27 supplement (Gibco Invitrogen, CA, USA), 0.25% GlutaMAX (Gibco Invitrogen, CA, USA), and 1% penicillin-streptomycin (PAN Biotech, Aidenbach, Germany). For immunocytochemistry investigation, cells were seeded on coverslips covered with poly-L-ornithine (0.1 mg/mL in borate solution) (PLO; Sigma-Aldrich, MO, USA) in 24-well culture plates (Corning, Merck, Darmstadt, Germany) in a density of 150000 cells/well. The cells were maintained at 37°C with 5% CO_2_, monitored closely and a new medium was added every 3-4 days.

#### Lead exposure and aSyn species treatment of primary neuronal cultures

Lead (II) acetate trihydrate (Sigma-Aldrich Chemie, Germany) was first diluted to 1M concentration in bi-distilled water. The initial concentration was then diluted into several intermediate concentrations of 500µM (0.5mM), 1mM, 5mM and 10mM solutions that were later added to the media of the primary neuronal cultures to dilute these into 1:1000 concentration to the final concentrations of 500nM (0.5µM), 1, 5, and 10µM in the cultures. For permanent exposure, the intermediate lead solution was added on the day *in vitro* 5 (DIV5) and left in the neuronal culture until the day *in vitro* 21 (DIV21). For intermittent exposure, the intermediate lead solution was also added on the day *in vitro* 5 (DIV5), the lead solution was withdrawn on the day *in vitro* 7 (DIV7) by changing the media of the primary neuronal cultures, and then added again on the day *in vitro* 14 (DIV14) until DIV21. The cells were closely monitored throughout the entire lead exposure time under the Whitefield microscope.

On day *in vitro* 7 (DIV7), after the media change in the cells, the cells were treated with 100nM of the different aSyn species, namely Monomers and Preformed fibrils (PFFs). PBS was used as a vehicle and control of the treatment. For Monomers solution preparation, the aliquots were thawed in ice and centrifuged at 14000g at 4°C for 25 min. The supernatant was taken out and diluted in sterile Dulbecco’s phosphate-buffered saline (DPBS, PAN Biotech, Germany) and added to the cells to a final concentration of 100nM. The PFFs were first thawed at room temperature, diluted in DPBS, and then sonicated for 1 min in the bath-type ultrasonicator (Branson 1510, Danbury, CT). The fibrils were then added to the cells at a final concentration of 100nM.

#### H4 cell culture and transfection

Human neuroglioma cells (H4) were maintained in Opti-MEM I Reduced Serum Medium (Life Technologies-Gibco, Carlsbad CA, USA) supplemented with 10% Foetal Bovine Serum Gold (FBS) (PAA, Cölbe, Germany) and 1% Penicillin-Streptomycin (PAN, Aidenbach, Germany). The cells were grown at 37°C with a 5% CO_2_ environment. Twenty-four hours prior to transfection, approximately 80000 H4 cells/well were plated in a 12-well plate (Costar, Corning, New York, USA) in 1mL Optimem medium. Six hours prior to transfection medium was replaced with a fresh one.

Transfection was performed using FuGENE® HD Transfection Reagent, as described by the manufacturer. Briefly, the transfection reagent was removed from the 4°C and left to get to room temperature. The vial of FuGENE® HD Transfection Reagent was vortexed until no precipitate was visible. In a 2mL Eppendorf tube, FuGENE® HD Transfection Reagent was added directly in a specific calculated volume in Optimem medium without adds. The ratio of 1 (equal amounts of the plasmids encoding SynT with Synphilin-1-V5) :3 (FuGENE® HD Transfection Reagent). The final prepared mix was calculated as 100uL to be added to the cells in the wells. For example, for 1 well, 1ug of SynT plasmid + 1ug Synphilin-1-V5 plasmid (total of 2ug) + 6uL of FuGENE® solution + 100uL Optimem without adds media was prepared. The mix was incubated for 30 min and added dropwise to the cells while the plate was gently rocked.

#### Lead exposure of H4 cells

As for the primary neuronal cultures, lead (II) acetate trihydrate (Sigma-Aldrich Chemie, Germany) was first diluted to 1M concentration in bi-distilled water. This initial concentration was then diluted to the intermediate concentrations of 50mM and 100mM solutions that were used for the experiments in a 1:1000 dilution to final concentrations of 50 and 100uM in the cultures.

For permanent exposure, 24 hours post-transfection, the lead solution was added and left on the plate for 24 hours. For intermittent lead exposure, the cells were exposed to lead after 24 hours of transfection, the lead was removed after 8 hours of exposure (the medium was aspirated and new Optimem medium added) for 12 hours and the lead was added again for another 4 hours. A total of 24-hour exposure protocol was applied in an intermittent paradigm.

### Immunocytochemistry (ICC)

At the end of the lead exposure protocol, primary whole brain neuronal cultures and H4 cells were washed with 1x DPBS (PAN Biotech, Aidenbach, Germany) and fixed with 4% of paraformaldehyde solution (PFA) for 20 min (home-made) at room temperature (RT). Cells were then washed 3 times with DPBS 1x solution (PAN Biotech, Aidenbach, Germany) and permeabilized with 0.1% Triton X-100 (Sigma-Aldrich, MO, USA) for 10 min at RT, and then incubated with 3% BSA in PBS (NZYTech) blocking solution for 1h at RT. Incubation with primary antibodies (1:1000) was performed overnight at 4°C [primary neuronal cultures: MAP2 (mouse, ab11267, abcam), phosphorylated alpha synuclein S129 – paSyn S129 (rabbit, ab51253, abcam) and H4 cells: alpha synuclein (mouse, 610787, BD Biosciences)].

Subsequently, the cells were washed three times with PBS 1x solution (PAN Biotech, Aidenbach, Germany) and then incubated with fluorescence conjugated secondary antibodies (1:1000) for 2h at RT [primary neuronal cultures: Alexa Fluor 488 goat anti-rabbit (A11008, Invitrogen) and Alexa Fluor 568 goat anti-mouse (A-11004, ThermoFisher) and H4 cells: Alexa Fluor 488 goat anti-mouse (A11029, Invitrogen)]. Finally, cells were stained with DAPI (Carl Roth, Karlsruhe, Germany) (1:10000 in PBS 1x) for 10 minutes, and the coverslips mounted in SuperFrost® Microscope Slides treated with Mowiol (Calbiochem, San Diego, CA) dried and stored at room temperature until further visualization and analysis.

### Imaging and analysis

Z-stack images of the primary neuronal cells were taken with a confocal point-scanning microscope (Zeiss LSM 900 with Airyscan 2, Carl Zeiss AG, Oberkochen, Germany) with a 10x objective (Objective Plan-Apochromat 10x/0.45 M27) with tile scan, 20x objective (Objective Plan-Apochromat 20x/0.8) and 63x objective (Objective Plan-Apochromat 63x/1.4 Oil DIC M27) for zoom in images of the neurons for fibril size analysis. For H4 cells, z-stack images were acquired in the same confocal microscope using the 63x objective with specific definitions for each staining.

Zen Microscopy Software (3.4 version, Carl Zeiss AG, Germany) was used for all the imaging experiments and the final images were post-analyzed and quantified using Fiji open-source software (Schindelin *et al*., 2012).

### Primary neuronal cell image analysis

For quantification of the percentage of neuronal cells with PFFs (paSyn S129-positive cells), the 10x tile images were selected and a 3D object counter plugin in Fiji was used to first count the MAP2-stained neuronal cells and afterwards count the number of paSyn S129-positive cells. The following equation was applied for calculating the percentage of cells: (paSyn S129-positive cells/MAP2-positive neurons) x 100. To quantify the size of the fibrils, individual fibrils (paSyn S129-positive structures) were selected inside the cells and their area was measured using Measure analysis of Fiji in the 63x images. To represent that these fibrils are inside the neurons, we are showing the orthogonal projection images.

### H4 cell image analysis

To quantify the number of aggregates and the size of the aggregates (area of the inclusions) inside the cells, the aSyn channel was selected and the images were thresholded. The images were then analyzed using the Analyze particle plugin from Fiji open-source software.

### In vivo studies

#### Lead exposure animal model

The lead exposure models were developed as described previously (Guimarães *et al*., 2012; Geraldes *et al*., 2016). Seven-day-pregnant Wistar rats (Charles River Laboratories, Chatillon-sur-Chalaronne, France) were separated into Pb-treated and control groups. The tap drinking water in the Pb-treated group was replaced with a 0.2% (p/v) lead (II) acetate solution (Acros Organics, New Jersey, NJ, USA) dissolved in deionized water.

The pups, after weaning at 21 days, were divided into 6 groups according to the type of exposure and brain injection. The 0.2% lead acetate solution was given to the lead exposed groups: intermittent groups (IntPb, n=48) – lead exposure until 12 weeks of age, no exposure (tap water) until 20 weeks, and second exposure from 20 to 28 weeks of age; permanent groups (PerPb, n=42) – lead solution in the animal diet until 28 weeks of age. Tap water was given to the age-matched control groups (Ctrl, n=38).

To offer a full functional and morphological evaluation, all animals were exposed to the same experimental methodology. The experimental methodology complied with European and national animal welfare regulations and was authorized by the Academic Medical Centre of Lisbon, Portugal (ref. 411/16).

#### Intrastriatal stereotaxic injections

Immediately before injection, recombinant aSyn PFF from the same batch as used for the *in vitro* studies, were thawed, diluted in sterile PBS solution, and sonicated for 1 minute in a bath-type ultrasonicator (Branson, Danbury, CT). Animals of 12 months of age were anaesthetized with a mixture of Ketamine/Dexmedetomidine, 0.2 ml/100g in weight, administered intraperitoneally and the withdrawal reflex was checked. Temperature was maintained through a homoeothermic blanket (Harvard Apparatus, Cambourne, UK).. When the withdrawal was not present, the animals were placed into a stereotaxic device (Kopf Instruments, Tujunga, CA, USA) and the head of the animal was fixed with ear bars from the device. Recombinant aSyn PFF solution or sterile PBS was injected into the right striatum at one site (8µg protein; AP +1.6, ML +2.4, DV−4.2 from the skull) at a rate of 0.5ul per minute using a microinjection pump (World Precision Instruments, Inc., Florida, USA). After each injection, the needle (10µL, 33-gauge Hamilton micro syringe (Hamilton Company, Nevada, USA)) was left in place for 2 min and then slowly withdrawn. For control rats, the same volume of vehicle solution (PBS) was injected. The skin of the rat was sutured with Silkam® 4/0 silk sutures (B-Braun, Melsungen, Germany) after the syringe was removed, and Bepanthene® Plus (Bayer, Leverkusen, Germany) was put on top of the sutures. Individual animals were kept in recovery cages with heating pads until they regained ambulatory ability. At the end of the stereotaxic surgery protocol, the animals were divided into 6 experimental groups according to the lead exposure and type of injection: Ctrl PBS, Ctrl aSyn; IntPb PBS, IntPb aSyn, PerPb PBS and PerPb aSyn.

### Behavioral Evaluation

The animals were subjected to a battery of standard behavioral tests two weeks prior to the functional evaluation (at 26 weeks of age). Behavioral evaluations were performed to assess (i) motor function and coordination with rotarod (Deacon, 2013; Laurent *et al*., 2016; Creed & Goldberg, 2018), (ii) anxiety-like behavior using elevated plus maze (Naolapo *et al*., 2012), (iii) locomotor and exploratory behavior with open filed test (Rogério dos Santos Alves; Alex Soares de Souza, 2014), and (iv) episodic long-term memory via novel object recognition test (Antunes & Biala, 2012). During the experimental days, animals were kept in the behavior testing room for at least an hour before the testing session began.

All behavioral experiments were conducted between the hours of 8 a.m. and 6 p.m. in a quiet room with low lighting, and all animals went through a four-day handling period (Schmitt & Hiemke, 1998; Costa *et al*., 2012) for researcher and testing room habituation and bias reduction. Between animals, all behavioral equipment was cleaned with 70% ethanol. All tests were videotaped using a UV camera (Chacon, Wavre, Belgium), and the movies were analyzed using ANY-maze software (Stoelting Co., Wood Dale, IL, USA).

#### Rotarod test

Motor function and coordination were tested on an accelerating rotarod (MED Associates Inc., St Albans, VT, USA) using the protocol described elsewhere (Deacon, 2013; Laurent *et al*., 2016; Creed & Goldberg, 2018; Lubrich *et al*., 2022). Rats were put on the rod with their heads pointing in the opposite direction of the rotation, forcing the animal to go forward to retain its equilibrium. Rats were first trained at a steady pace (7 r.p.m. for 5 min) before beginning three test trials (at least 30 min intertrial interval). Animals had to balance on a revolving rod that accelerated from 4 to 40 r.p.m. in 5 minutes during these trials. The time it took for the rats to fall off the revolving rod was timed to a maximum of 5 minutes and the r.p.m was also calculated.

#### Open-Field Exploration Test

Taking advantage of the curious nature of the rodents, the open field test (OFT) was used for exploration of a new environment and general locomotion (Buccafusco, 2001). The OFT apparatus is composed of a square black box (67 x 67 x 57 cm in height) that has been "virtually" split into three concentric squares: (1) the periphery zone (near the walls), (2) the intermediate zone, and (3) the center. The animals were left in the maze to explore freely for 5 minutes, which is generally enough time to assess the specified parameters. We computed the total travelled distance and the average velocity of the animals as the main parameters of the test (Moreira *et al*., 2001; Ramos, 2008; Rogério dos Santos Alves; Alex Soares de Souza, 2014; Shvachiy *et al*., 2018, 2020b).

#### Novel Object Recognition Test

Novel object recognition test (NOR) with a 24-hour retention interval was used to evaluate the episodic long-term memory by the same protocol as described previously using the OFT arena (Antunes & Biala, 2012; Shvachiy *et al*., 2018, 2020b, 2022). Briefly, the objects used (clear and brown glass shapes) were randomized and their position in relation to the other objects was changed to use each object as a source of familiarity or novelty.

The test consists of three stages: habituation, training, and testing. During habituation (three days), animals were left to explore the apparatus freely for 15 min. On the fourth day, they were exposed to two familiar (F) items, for 5 minutes. On the fifth day, the animals were exposed to two objects for 5 minutes, one previously encountered object (F) and one novel object (N) (Antunes & Biala, 2012). The testing day was recorded and analyzed using ANY-maze® software employing 3-point analysis (head, torso, and tail of the animal), and only the data from the head point analysis were relevant for object exploration. The amount of time animals spent around each object during the testing stage was used to quantify exploratory behavior. The number of approaches that involved smelling, rearing towards, or touching the item was tallied. Exploration did not include sitting rearward to the item or passing in front of it without pointing the nose in the object’s direction(Antunes & Biala, 2012). Exploration time was measured as follows: ET (%) = (time exploring the object/overall exploring time) × 100.

The novelty index was computed as follows:

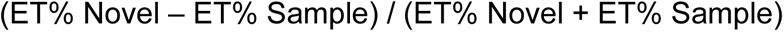

This index has a range of −1 to 1, with negative values representing the lack of discrimination between the new and familiar items (i.e., spending more time exploring the sample object or equal time exploring both items) and positive values representing more exploration of the novel object (Mouro *et al*., 2017; Shvachiy *et al*., 2018, 2020b).

### Metabolic Evaluation

Rats were housed in metabolic cages for 24 hours before the acute experiment at 28 weeks to examine body weight, food and drink consumption, and urine and feces production.

### Functional Evaluation

#### Physiological and autonomic evaluation

After behavioral and metabolic evaluation, animals were anaesthetized with sodium pentobarbital (60 mg/kg, intraperitoneal injection). Anesthesia levels were maintained using a 20% solution (v/v) of the same anesthetic after assessing the withdrawal response. A homoeothermic blanket (Harvard Apparatus, Cambourne, UK) was used to keep the animal’s temperature stable. To measure tracheal pressure, the trachea was cannulated below the larynx. The femoral artery and vein were cannulated to monitor blood pressure (BP) and inject saline and drugs, respectively. The electrocardiogram (ECG) was assessed using subcutaneous electrodes in three limbs, and the heart rate was calculated using the ECG data (Neurology, Digitimer, Welwyn Garden City, UK).

The right carotid artery was catheterized, and chemoreceptors were stimulated with lobeline (0.2 mL, 25 g/mL, Sigma, St. Louis, MO, USA) (Geraldes *et al*., 2016). Phenylephrine injection (0.2 mL, 25 g/mL, Sigma, St. Louis, MO, USA) in the femoral vein was used to stimulate the baroreflexes (Geraldes *et al*., 2016; Shvachiy *et al*., 2018, 2020b).

Blood lead levels (BLL) were determined from the venous blood using an atomic absorption spectrophotometer (Shimadzu, Model no. AA 7000, Kyoto, Japan). At the end of the above-mentioned acute experience, the animal was sacrificed with an overdose of anesthetic.

#### Data Acquisition and Analysis

Continuous recordings of blood pressure (BP), ECG, heart rate (HR), and respiratory frequency (RF) were performed (PowerLab, ADInstruments, Colorado Springs, CO, USA) and acquired, amplified, and filtered at 1 kHz (Neurology, Digitimer, Welwyn Garden City, UK; PowerLab, ADInstruments, Colorado Springs, CO, USA). A baseline recording of 10 minutes was taken for basal autonomic assessment. Each stimulus was separated by at least 3 minutes.

#### Baro- and Chemoreceptor Reflex Analysis

The autonomic reflexes analysis was focused on assessing baro- and chemoreceptor responses. The baroreceptor reflex gain (BRG) was calculated after phenylephrine provocation by measuring ΔHR/ΔBP (bpm/mmHg).

The chemoreflex response was estimated using respiratory frequency (RF) measured from tracheal pressure before and after lobeline stimulation: ΔRF = RFstimulation - RFbasal.

### Immunohistochemistry (IHC)

After sacrifice, animals underwent transcardial perfusion with fresh PBS 1x solution (TH Geyer, Germany), the brain was removed and transferred into paraformaldehyde (PFA 4%, Carl Roth, Germany) solution for post-fixation for 72h. The brains were then immersed in increasing concentrations of sucrose (15% and 30%, Merck, Germany in PBS 1x with sodium azide, Sigma, Germany) and kept at 4°C for later analysis.

To evaluate the astrocytic, alpha-synuclein, phosphorylated alpha-synuclein and dopaminergic neurons changes, sagittal free-floating sections (30μm) were cut around the region of hippocampus [Lateral 0.6 – 2.04 using a cryostat (Leica CM 3050S, Germany)] and, subsequently stained as reported previously (Chegão *et al*., n.d.; Shvachiy *et al*., 2022). Succinctly, free-floating sections were washed with TBS 1x (PanReac/AppliChem, Germany), permeabilized with 3% Triton X 100 solution for 15min (Carl Roth, Germany) and blocked with 5% Goat Serum (BioWest, France) and 1% Bovine serum (VWR, USA) for 1 h. The freely floating tissue were then incubated with specific combinations of primary antibodies [astrocytes - glial fibrillary acidic protein - GFAP (chicken, ab4674, abcam, 1:500), alpha synuclein (mouse, 610787, BD Biosciences), phosphorylated alpha synuclein S129 – paSyn S129 (rabbit, SMC-600, StressMarq), Tyrosine Hydroxylase - TH (mouse, MAB5280, Milipore) and Tyrosine Hydroxylase – TH (rabbit, AB152, Milipore)] overnight at 4°C on a shaker.

Sections were then washed with TBS 1x and incubated with specific combinations of secondary antibodies [Alexa Fluor 488 donkey anti-mouse (A21202, ThermoFisher), Alexa Fluor 568 goat anti-rabbit (A11036, Invitrogen), Alexa Fluor 633 goat anti-chicken (A21103, Invitrogen), Alexa fluor 488 donkey anti-rabbit (A21206, Invitrogen) and Alexa Fluor 568 goat anti-mouse (A-11004, ThermoFisher)]. Lastly, nuclei were counter-stained with DAPI (4′,6-diamidino-2-phenylindole, 1:10000, Carl Roth, Germany). Sections were mounted in SuperFrost® Microscope Slides using Fluoromount-G mounting media (Invitrogen, Germany). Omission of the primary antibody resulted in no staining.

Z-stack images of the dentate gyrus region of the hippocampus, striatum and *substantia nigra* were taken with a confocal point-scanning microscope (Zeiss LSM 900 with Airyscan 2, Carl Zeiss AG, Oberkochen, Germany) with a 20x objective (Objective Plan-Apochromat 20x/0.8) and tile scan to image the specific region of interest and 63x objective (Objective Plan-Apochromat 63x/1.4 Oil DIC M27) for zoom in images of the cells for morphological evaluation of the astrocytic cells. Zen Microscopy Software (3.4 version, Carl Zeiss AG, Germany) was used for all the imaging experiments and the final images were post-analyzed and quantified using Fiji open-source software (Schindelin *et al*., 2012).

For morphological evaluation of the astrocytes, relevant scientific articles were used for support (Sofroniew & Vinters, 2010; Hol & Pekny, 2015; Siracusa *et al*., 2019). For quantification of the percentage of GFAP-positive cells, the dentate gyrus region of the 20x tile images was selected and a 3D object counter plugin in Fiji was used to first count the DAPI-stained nuclei and afterwards count the number of GFAP-positive cells. The following equation was applied for calculating the percentage of cells: (GFAP-positive cells/DAPI nuclei) x 100. Fluorescence intensity inside the cells and their area were measured using Measure analysis of Fiji in the 63x images with cells individually selected for analysis. For evaluation of the fluorescence intensity, the selected region of the dentate gyrus and the *substantia nigra* and the whole image of the striatum, the images were measured with the measure analysis from Fiji open-source software.

### Statistical Analysis

If not otherwise specified, the data were presented as mean ± SD and represented the average of mean values across all animals/cells/biological repetitions. The normality distribution of continuous variables was assessed using the D’Agostino and Pearson normality test, and the homogeneity of variance was examined using Levene’s test. For data analysis among different groups, both in *in vitro* and *in vivo* studies encompassing behavioral, physiological, and molecular parameters, a one-way ANOVA was employed. Subsequently Tukey’s multiple comparisons test was applied for normally distributed data, and Kruskal-Wallis multiple comparison test for non-normally distributed values. Additionally, to compare the exploration time percentage between objects in the NOR test for each group, a student t-test for paired observations was used. The statistical analysis was performed using GraphPad Prism 9 (GraphPad Software Inc., USA). Statistical significance was defined as p < 0.05.

## Results

### Lead exposure increases aSyn inclusion formation in H4 cells

First, to study the effect of lead exposure on aSyn inclusion formation, we used an established model of aggregation based on the co-expression of a modified form of aSyn (SynT) and synphilin-1(McLean *et al*., 2001; Lázaro *et al*., 2017). Interestingly, we found that both types of lead treatment profiles induced an increase in the number of inclusions per cell (Fig. 1A-B) but not in the average size of the inclusions (Fig. 1C).

**Figure 1.**
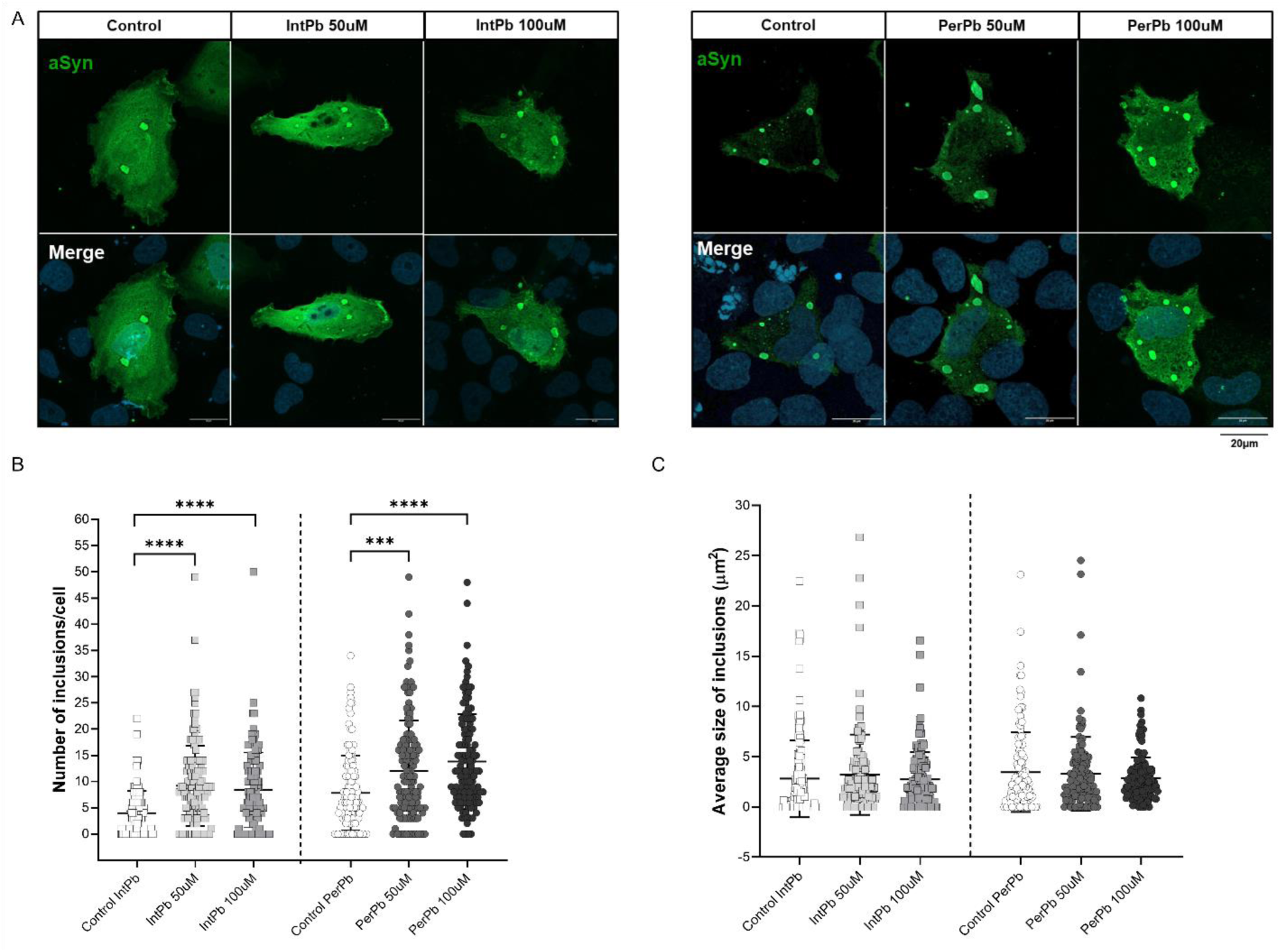
Effect of lead on aSyn inclusion formation in H4 cells. A. Representative images of cells treated with lead and immunostained with an antibody against aSyn. B. Quantification of aSyn-positive inclusions inside the cells. C. Histogram of the average size of the inclusions. Images were acquired using a confocal point scanning microscope (Zeiss LSM 900 with Airyscan2) with a 63× objective. Scale bar is 20 µm. Values are the mean ± SD (n=123-163 cells) with 3 independent experiments for each evaluated group. Symbols refer to statistically significant differences inter groups (Ctrl vs IntPb/PerPb - *p < 0.05, **p < 0.01, ***p < 0.001, ****p < 0.0001); one-way ANOVA, Kruskal-Wallis multiple comparison test for non-normal values.

### Lead exposure increases the number and size of inclusions in aSyn PFF-treated primary neuronal cultures

In order to assess the effect of lead in a neuronal cell model, we investigated the effect of lead in the generation of inclusion in primary neuronal cultures by the addition of pre-formed fibrils (PFFs). First, we confirmed the formation of inclusions in the cells by the addition of the different aSyn species, namely monomers and PFFs. As a control, we used the DPBS solution, being that the vehicle of the preparation of the species.

The orthogonal projections of the representative images of paSyn S129 staining for aggregates and MAP2 marker for neurons show that the PFFs treatment generated abundant aggregate formations in the cells in all groups. The orthogonal projection shows that these formations are inside the neurons (the green staining – paSyn S129 is enclosed in the red staining – MAP2, as observed in Fig. 2A) Curiously, we observed that neither PBS nor Mon lead to the formation of inclusions in the primary neuronal cells. Interestingly, we detected that the aggregates formed in the PFFs groups increase in number with the presence of lead with both intermittent and permanent lead exposure, with a significant increase in the percentage of neurons with inclusion in the PerPb 5µM group (Fig. 2C).

**Figure 2.**
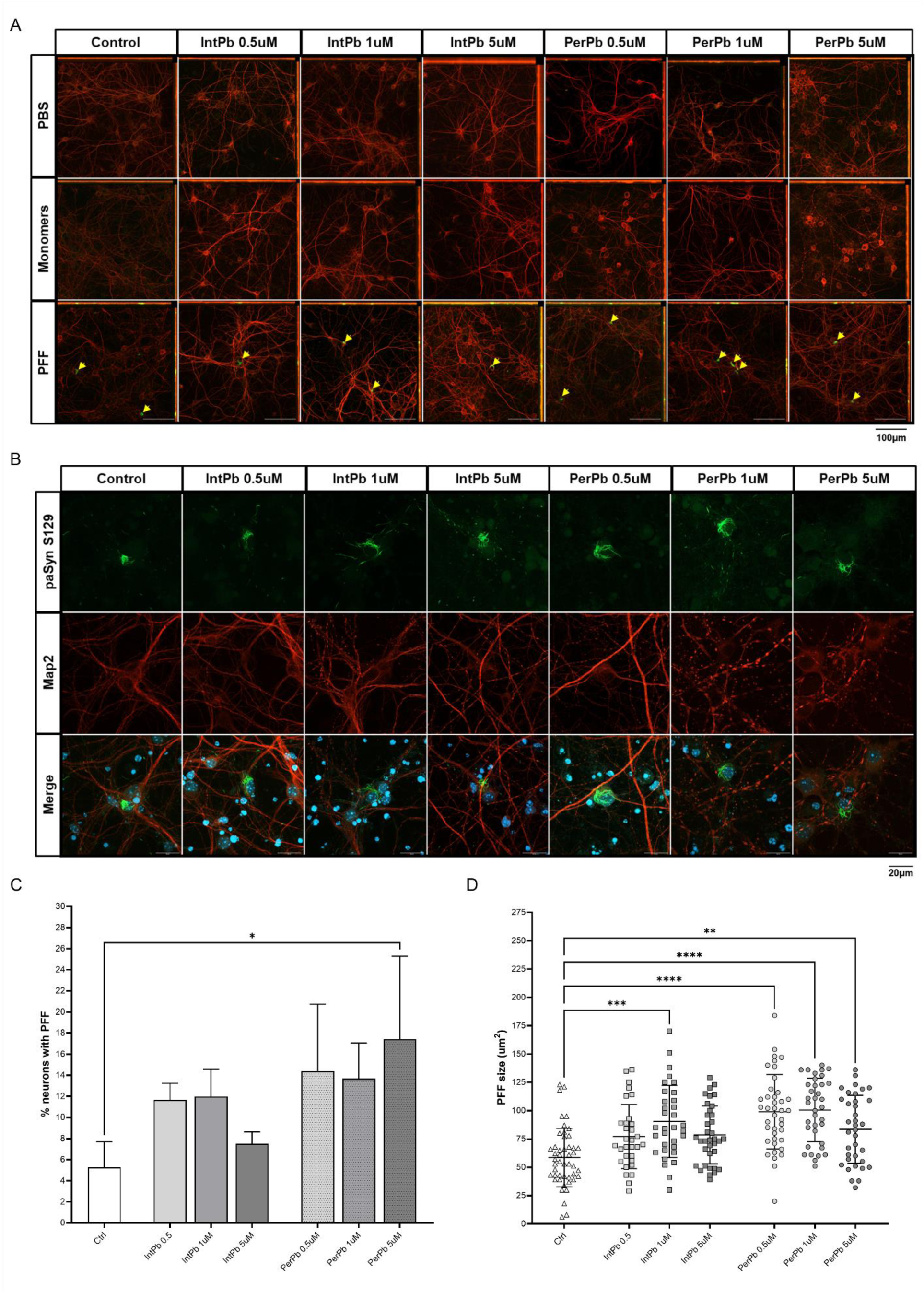
Effect of lead exposure on aSyn aggregation in primary neuronal culture. A. Representative images of the orthogonal projection images of neuronal cells treated with the different aSyn species, stained with paSyn antibody (1:1000), exposed to lead and controls. B. Representative images of the 63x magnification images of neuronal cells treated with the different aSyn species, stained with paSyn antibody (1:1000), exposed to lead and controls C. Histogram of the quantification of the number of neurons with paSyn-stained aggregates inside the cells of 10x magnification images. D. Histogram of the quantification of the size of the paSyn-stained aggregates inside the neurons. Images were acquired using a confocal point scanning microscope (Zeiss LSM 900 with Airyscan2) with 10x, 20x and 63x objectives. Scale bar is 100 µm and 20 µm for representative images. Values are the mean ± SD (n=3-5 independent experiments for each evaluated group and n=31-47 cells). The symbols denote statistically significant differences inter groups (Ctrl vs IntPb/PerPb - *p < 0.05, **p < 0.01, ***p < 0.001, ****p < 0.0001); one-way ANOVA, Tuckey’s multiple comparison test and Kruskal-Wallis multiple comparison test for non-normal values.

To evaluate the effect of lead on the size of aggregates in primary neurons, confocal images were taken and analysed. Lead exposure caused an increase in the size of the aggregates inside the cells (Fig. 2B). Likewise, the neuronal cells from IntPb 1 and 5µM and PerPb 1 and 5µM appear to be in stronger distress as the cells show axonal damage (intermittent MAP2 signal), decreased growth of the dendrites and death of the cell body. The quantifications of the images showed that lead exposure, regardless of the type, causes an increase in the size of the inclusions. However, only the IntPb 1uM from the intermittent groups shows a significant increase. The IntPb 0.5 and 5µM groups did not cause a significant increase in the size of the aSyn inclusions. As for the permanent exposure groups, all PerPb groups show a significant increase in the size of the inclusions (Fig.2D).

### Lead causes locomotor and exploration impairment in the aSyn PFF model without effect in coordination and balance injection

Next, to assess the effect of lead or aSyn PFF on motor performance of Wistar rats, we used the rotarod test. We observed no significant changes between the groups, regardless of the type of lead exposure or the injection, in the time spent on the rotarod (Fig. 3A). Therefore, no significant differences were observed in the rotations per minute (Fig. 3B).

**Figure 3.**
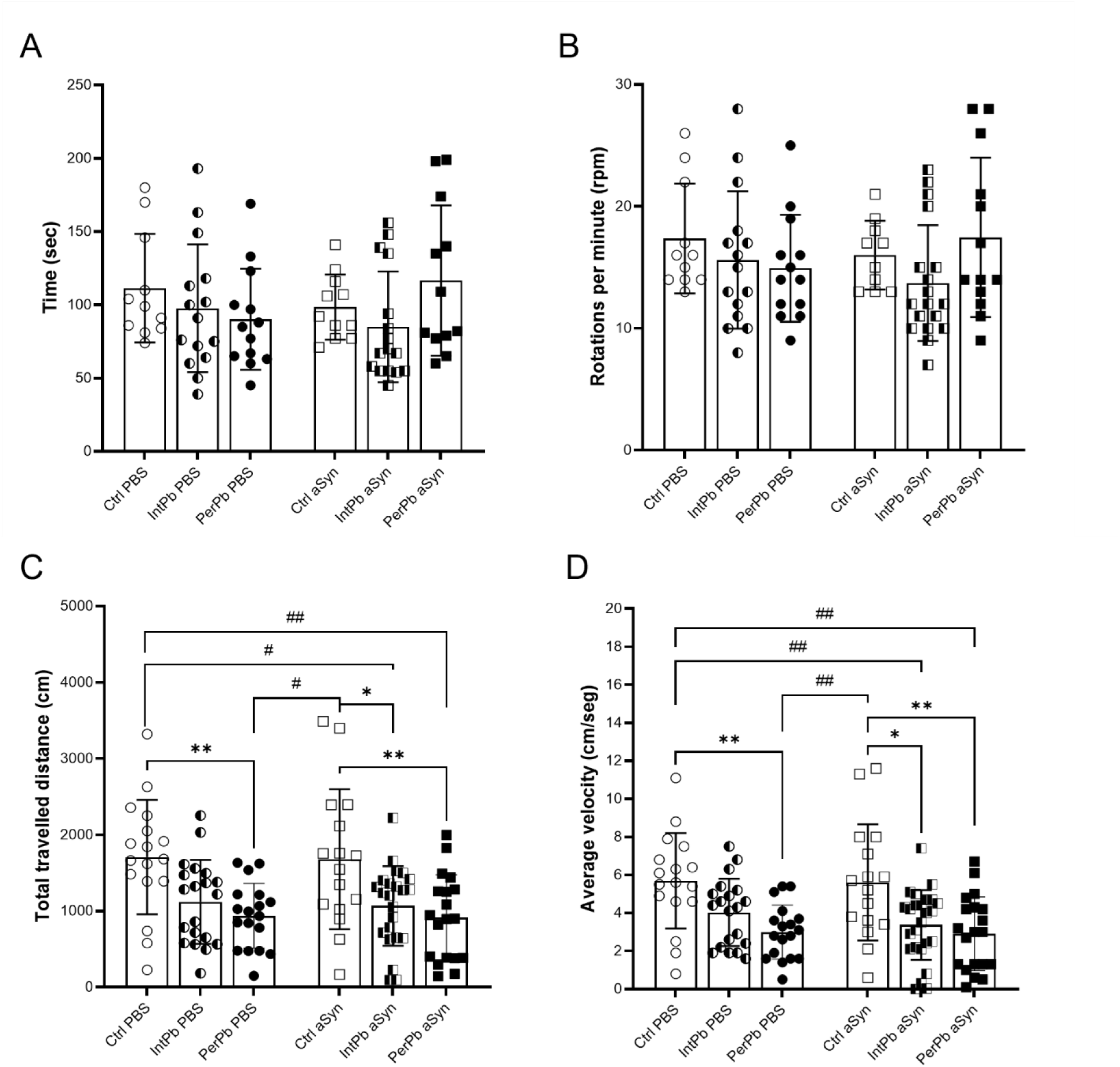
Effect of lead exposure and aSyn PFF injection in the motor coordination balance, locomotor and exploratory behaviour. A. Time spent in the rotarod apparatus or latency to fall. B. Rotations per minute on the rotarod apparatus. C. Total travelled distance in the open field test. D. Average velocity in the open field test. Values are the mean ± SD. The symbols denote statistically significant differences inter groups: Ctrl PBS vs IntPb PBS/PerPb PBS; Ctrl aSyn vs IntPb aSyn/PerPb aSyn - *p < 0.05, **p < 0.01, ***p < 0.001, ****p < 0.0001; Ctrl PBS vs IntPb aSyn/PerPb aSyn; one-way ANOVA, Tuckey’s multiple comparison test.

To evaluate the locomotor and exploratory behavior of the animals, the open field test was used. A significant decrease in total travelled distance was observed in PerPb PBS when compared to Ctrl PBS and when compared to Ctrl aSyn, and in both IntPb aSyn and PerPb aSyn when compared to Ctrl aSyn and to Ctrl PBS with significant changes between IntPb PBS with other groups (Fig. 3C).

A similar pattern of changes was observed in the average velocity. Namely, a significant decrease was observed in the PerPb PBS compared to Ctrl PBS and Ctrl aSyn as well as IntPb aSyn and PerPb aSyn showed a significant decrease when compared to Ctrl aSyn and Ctrl PBS without significant differences between IntPb PBS and other groups (Fig. 3D).

### Lead impairs episodic long-term memory alongside aSyn PFF injection

Next, we used the novel object recognition test to evaluate effects in episodic long-term memory with a retention interval of 24 hours. Regarding the exploration time, we observed that only the Ctrl PBS group shows a significant reduction in this parameter (Fig. 4A). No significant changes were observed in the other groups (Fig. 4A). Interestingly, when the novelty recognition index was calculated, we observed that both lead exposed groups, without aSyn, showed a significant decrease in this parameter (Fig. 4B). Similarly, the PerPb aSyn and Ctrl aSyn groups presented a significant decrease in the novelty recognition index when compared to Ctrl PBS. The IntPb aSyn group did not show significant differences when compared to the other groups (Fig. 4B).

**Figure 4.**
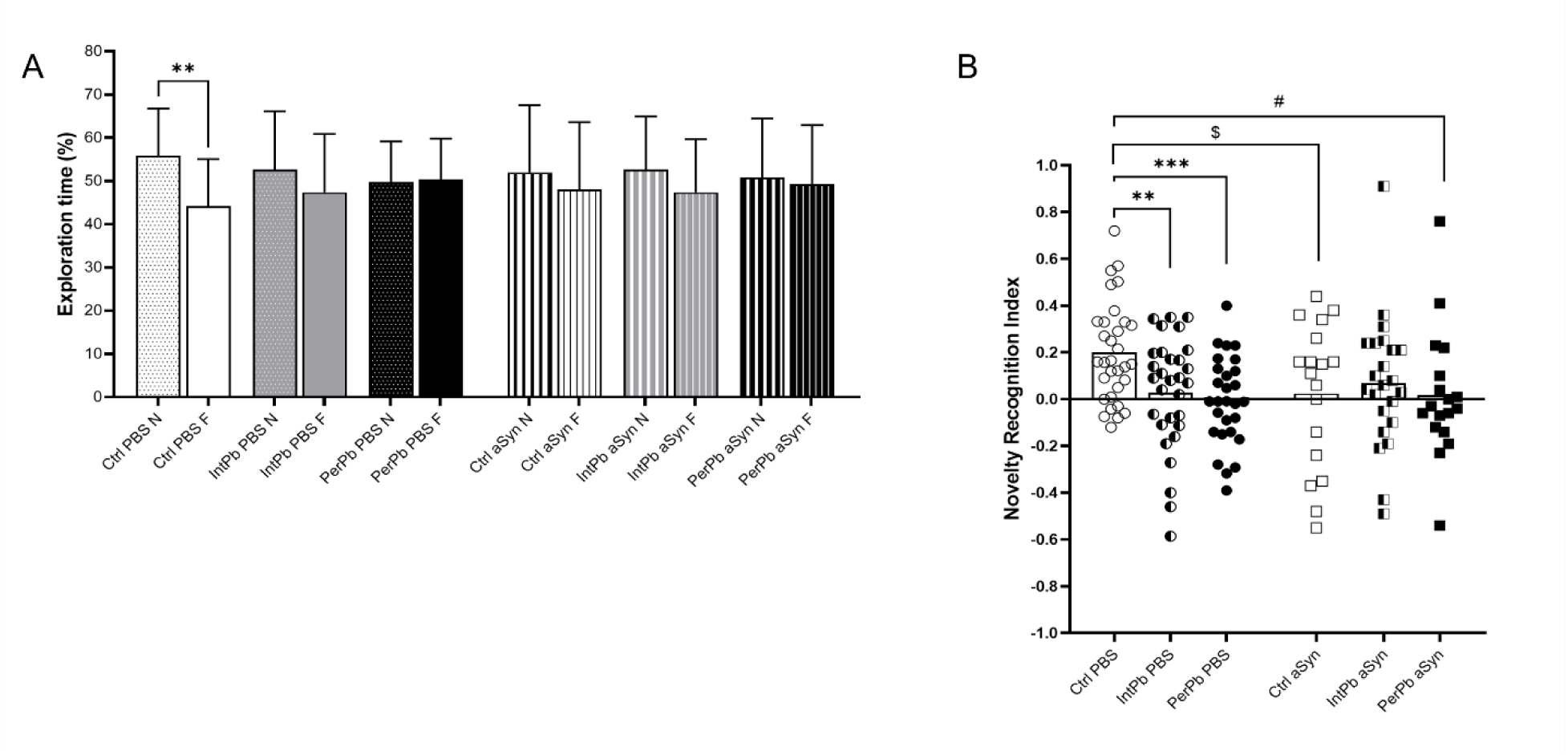
Effect of lead exposure and aSyn PFF injection in episodic long-term memory. A. Exploration time percentage of novel (N) and familiar (F) objects. B. Novelty recognition index was calculated. Values are the mean ± SD. The symbols denote statistically significant differences inter groups: Ctrl PBS vs IntPb PBS/PerPb PBS; Ctrl aSyn vs IntPb aSyn/PerPb aSyn - *p < 0.05, **p < 0.01, ***p < 0.001, ****p < 0.0001; Ctrl PBS vs IntPb aSyn/PerPb aSyn; Ctrl aSyn vs IntPb PBS/PerPb PBS - ^#^p < 0.05, ^##^p < 0.01, ^###^p < 0.001, ^####^p < 0.0001; Ctrl PBS vs Ctrl aSyn/ IntPb PBS vs IntPb aSyn/ PerPb PBS vs PerPb aSyn - ^$^p < 0.05, ^$$^p < 0.01, ^$$$^p < 0.001, ^$$$$^p < 0.0001; one-way ANOVA, Tuckey’s multiple comparison test; the differences between the objects (N and F) in the exploration time percentage was analyzed with Paired t-test.

### Lead increases blood pressure, respiratory frequency and chemoreceptor reflex sensitivity

During the acute surgery at 28 weeks of age, basal physiological parameters were evaluated 10 min before the stimulations of the autonomic reflexes. We observed that, regardless of the lead exposure and/or injection of aSyn, the animals showed a significant increase in the systolic, diastolic and, consequently, mean blood pressure, when compared to the Ctrl PBS group (Fig. 5A), but no changes in heart rate (Fig. 5B). A significant increase was observed in the respiratory frequency in the permanently lead-exposed animals, independently of aSyn injection, when compared to the Ctrl PBS group with no other significant changes in other groups (Fig. 5C).

**Figure 5.**
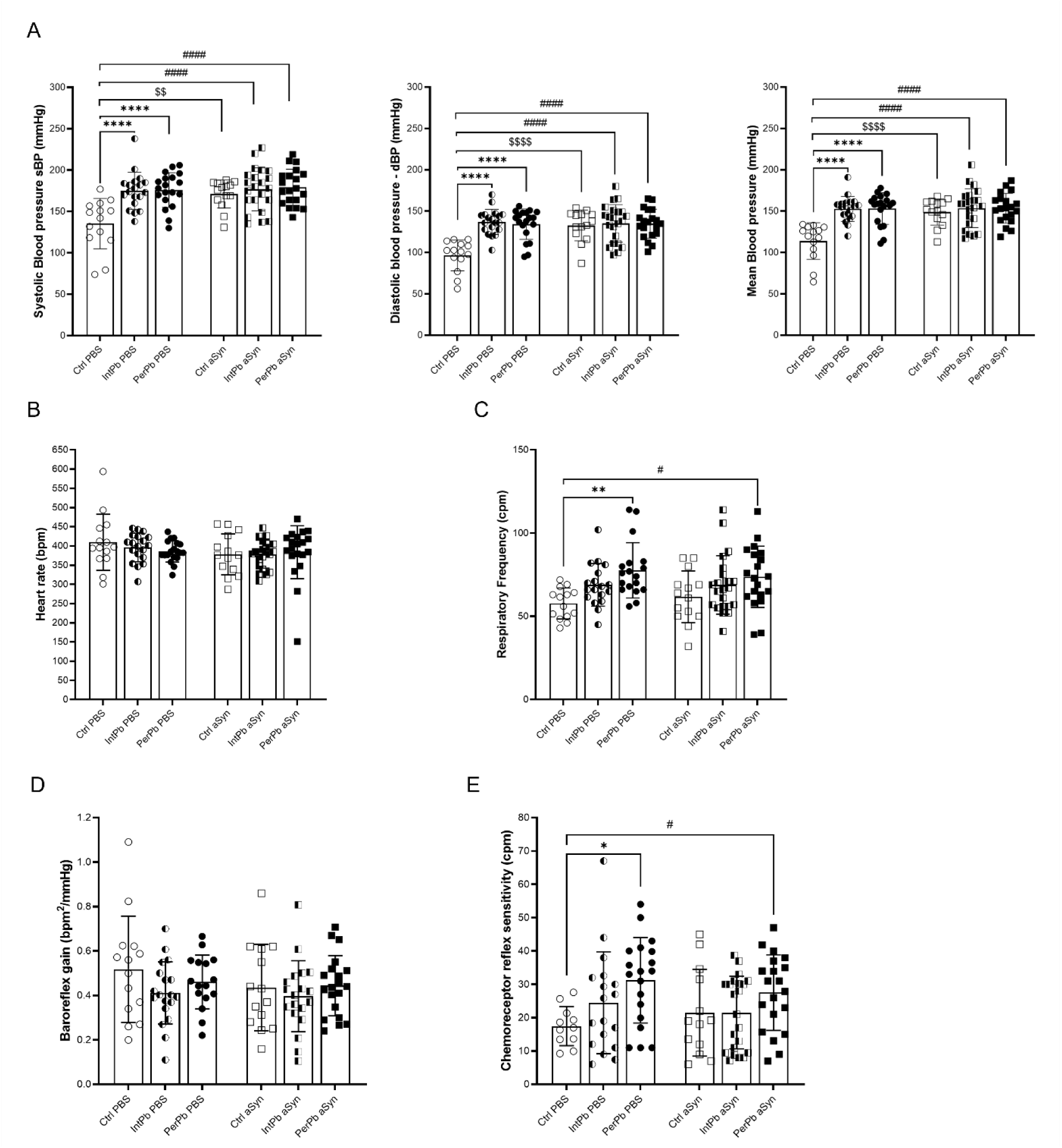
Effect of lead exposure and aSyn PFF injection in basal physiological parameter and autonomic reflexes. A. Systolic, diastolic, and mean blood pressure. B. Heart rate. C. Respiratory frequency. D. Baroreflex gain. E. Chemoreceptor reflex sensitivity. Values are the mean ± SD. The symbols denote statistically significant differences inter groups: Ctrl PBS vs IntPb PBS/PerPb PBS; Ctrl aSyn vs IntPb aSyn/PerPb aSyn - *p < 0.05, **p < 0.01, ***p < 0.001, ****p < 0.0001; Ctrl PBS vs IntPb aSyn/PerPb aSyn; Ctrl aSyn vs IntPb PBS/PerPb PBS - ^#^p < 0.05, ^##^p < 0.01, ^###^p < 0.001, ^####^p < 0.0001; Ctrl PBS vs Ctrl aSyn/ IntPb PBS vs IntPb aSyn/ PerPb PBS vs PerPb aSyn - ^$^p < 0.05, ^$$^p < 0.01, ^$$$^p < 0.001, ^$$$$^p < 0.0001; one-way ANOVA, Tuckey’s multiple comparison test.

After evaluating the basal physiological parameters, the autonomic reflexes were pharmacologically stimulated. We observed that, even though there is a slight decrease in the baroreflex gain, no significant differences were present in all the groups, regardless of lead exposure or injection (Fig. 5D). Interestingly, the PerPb group, regardless of the injection, shows a significant increase in the chemoreceptor reflex sensitivity, when compared to the Ctrl PBS group with no other significant changes in this parameter (Fig. 5E).

### Lead increases BLL and increases water intake and urine production in a model of aSyn pathology

The metabolic evaluation was performed for 24 hours in metabolic cages after the behavioral testing at 28 weeks of age. We observed that the intermittent lead exposure group injected with aSyn (IntPb aSyn) animals had a strong decrease in the water intake when compared to both Ctrl aSyn and Ctrl PBS groups. Interestingly, the IntPb PBS and PerPb PBS groups showed a significant decrease in urine production when compared to the Ctrl aSyn group. The IntPb aSyn group, similarly to the water intake, showed a significant decrease in urine production when compared to the Ctrl aSyn group (Fig. 6B). Regarding blood lead levels, we observed that both lead exposures resulted in a significant increase in the BLL, with PerPb groups showing a higher level than IntPb (Fig. 6A).

**Figure 6.**
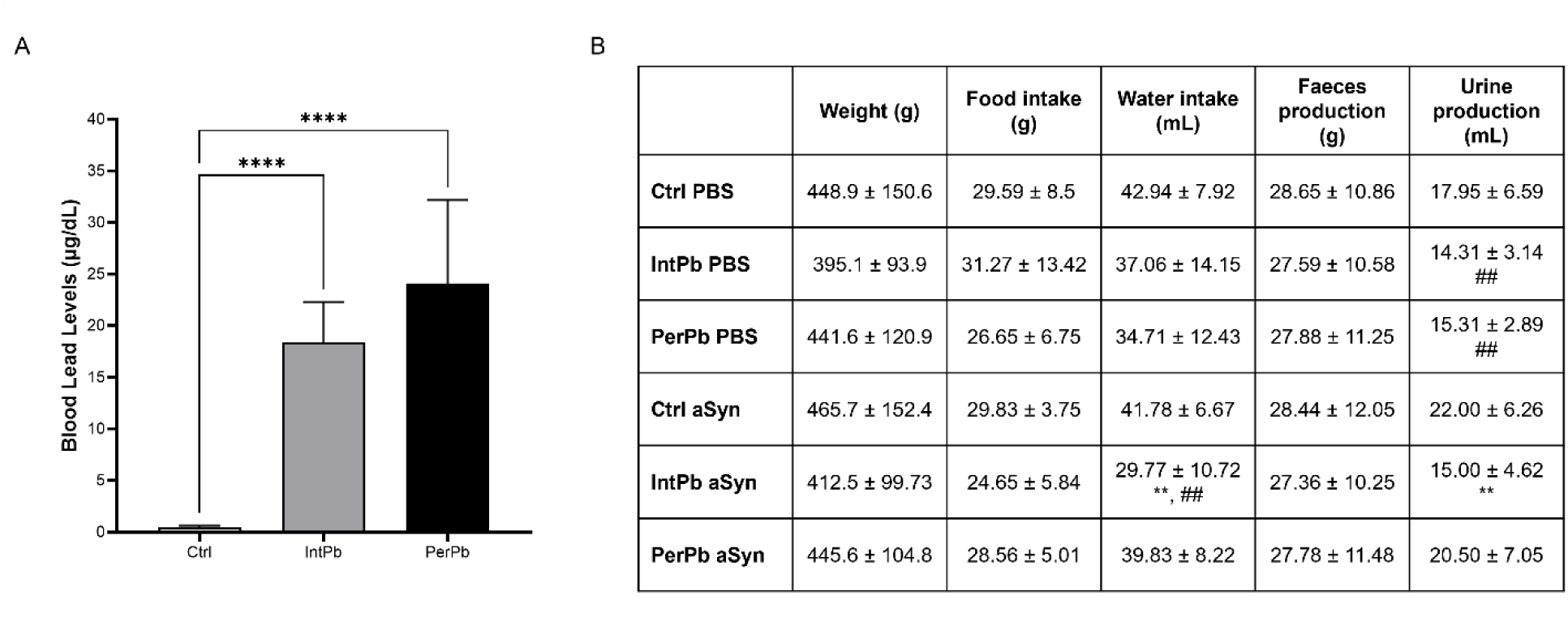
Effect of lead exposure and aSyn PFF injection in the metabolic parameters. A. Histogram of the blood lead levels evaluated by atomic spectrophotometry. B. Table with all data for the metabolic evaluation: weight, food and water intake, faeces, and urine production. Values are the mean ± SD (n=5-7 for blood lead levels; n=16-20 for metabolic data). The symbols denote statistically significant differences inter groups (Ctrl PBS vs IntPb PBS/PerPb PBS; Ctrl aSyn vs IntPb aSyn/PerPb aSyn - *p < 0.05, **p < 0.01, ***p < 0.001, ****p < 0.0001; Ctrl PBS vs IntPb aSyn/PerPb aSyn; Ctrl aSyn vs IntPb PBS/PerPb PBS - ^#^p < 0.05, ^##^p < 0.01, ^###^p < 0.001, ^####^p < 0.0001); one-way ANOVA, Tuckey’s multiple comparison test.

### Lead induces astrocytic atrophy alongside a reduction in aSyn pathology in the hippocampus

The morphological evaluation showed that lead-exposed groups without aSyn – IntPb PBS and PerPb PBS – showed a significant increase in the activation of the astrocytic cells, with hypertrophy of cellular processes and GFAP upregulation with a higher density of the astrocytic cells, confirming that astrocytes were in a reactive state (a hallmark of pathology). The same morphological changes were observed in the Ctrl aSyn group. IntPb aSyn and PerPb aSyn animals did not exhibit morphological alterations and were comparable to the Ctrl PBS group (Fig.7A). Our quantitative analysis confirmed the morphological alterations observed. Specifically, we observed a significant increase in the percentage of GFAP-positive cells in both lead-exposed groups (without aSyn injection) when compared to the Ctrl PBS group. Interestingly, the lead exposed groups with aSyn showed a decrease when compared to Ctrl aSyn. Significant differences were also observed between IntPb PBS and IntPb aSyn, as well as in PerPb PBS and PerPb aSyn (p<0.0001) with a significant decrease observed in the aSyn injected groups (Fig.7B). We observed differences in fluorescence intensity and in GFP-positive cells (Fig. 7C-E). Altogether, these findings suggest aSyn may modify astrocytic responses, in a cross talk that has been reported by others (Forloni, 2023; Li *et al*., 2024).

**Figure 7.**
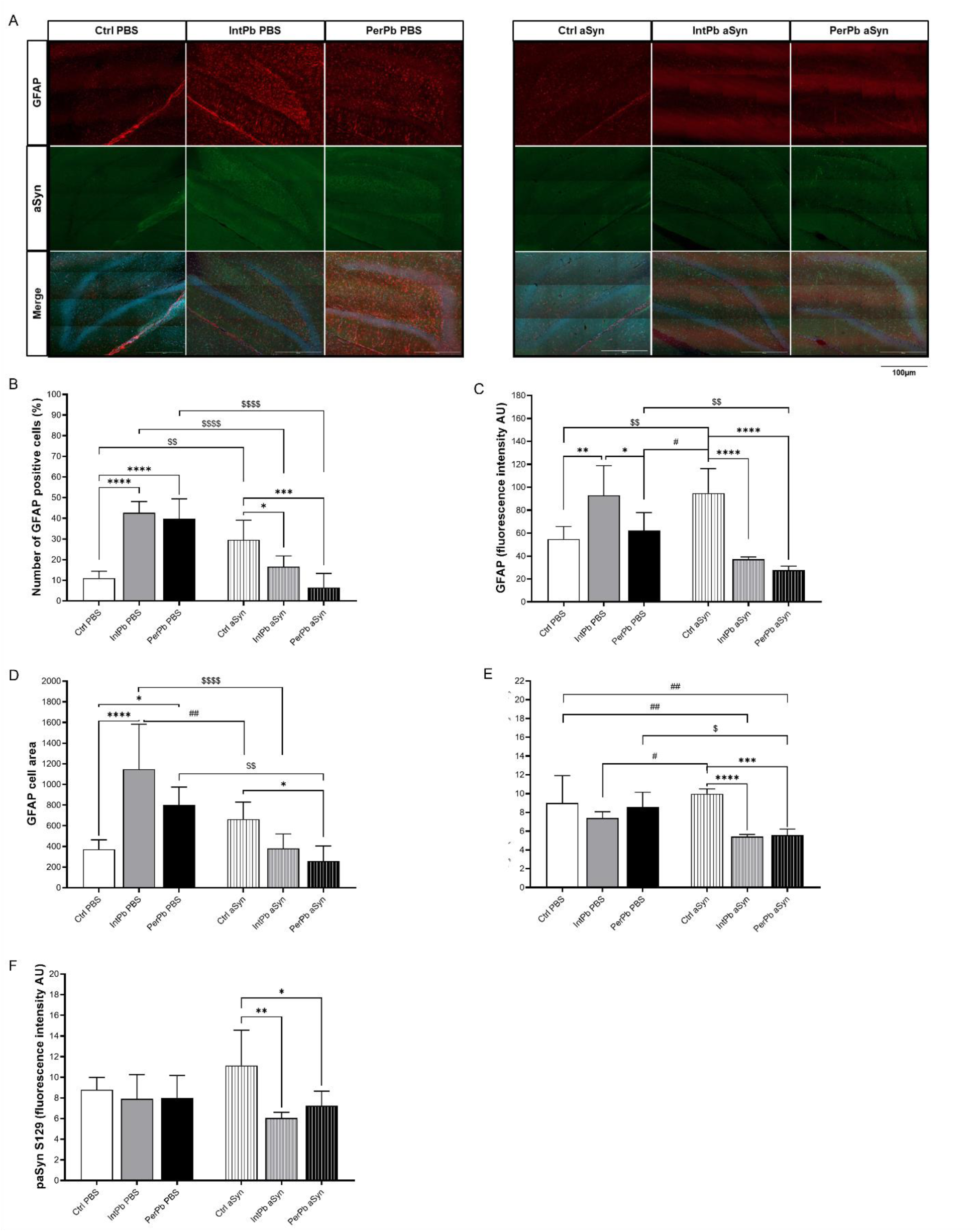
Effect of lead and aSyn PFF injection on astrocytic activation, aSyn and paSyn S129 levels evaluated in dentate gyrus of the hippocampus. A. Representative images of the GFAP (1:1000)- stained astrocytes and aSynuclein (1:200). B. Histogram of the percentage of GFAP-positive cells’ quantification. C. Histogram of GFAP fluorescence intensity quantification. D. Histogram of GFAP cell area quantification. E. Histogram of aSyn fluorescence intensity quantification. F. Histogram of paSyn S129 fluorescence intensity quantification. Images were acquired on a confocal point scanning microscope, (Zeiss LSM 900 with Airyscan 2), with a 20x objective for whole dentate gyrus imaging and a 63x objective for zooming in on the astrocytic cells. The scale bar is 100 µm or 50 µm for stained images. Values are the mean ± SD for n=3-8 for all groups. The symbols denote statistically significant differences inter groups: Ctrl PBS vs IntPb PBS/PerPb PBS; Ctrl aSyn vs IntPb aSyn/PerPb aSyn - *p < 0.05, **p < 0.01, ***p < 0.001, ****p < 0.0001; Ctrl PBS vs IntPb aSyn/PerPb aSyn; Ctrl aSyn vs IntPb PBS/PerPb PBS - ^#^p < 0.05, ^##^p < 0.01, ^###^p < 0.001, ^####^p < 0.0001; Ctrl PBS vs Ctrl aSyn/ IntPb PBS vs IntPb aSyn/ PerPb PBS vs PerPb aSyn - ^$^p < 0.05, ^$$^p < 0.01, ^$$$^p < 0.001, ^$$$$^p < 0.0001; one-way ANOVA, Tuckey’s multiple comparison test.

### Lead reduces dopaminergic neurons and aSyn levels in the substantia nigra and striatum

We observed a reduction in TH-positive cells in the intermittent lead exposure groups, and a slight increase in aSyn levels with the presence of lead or aSyn (Fig. 8A). The quantification of the images of the substantia nigra showed that the intermittent lead exposure, independent of the injection (PBS or aSyn) caused a significant decrease in the TH-positive cells when compared to the Ctrl PBS group (Fig. 8C). The quantification of the aSyn signal in the substantia nigra and striatum showed that, even though there is a minor increase in the fluorescence intensity, it is not significant in all groups, regardless of the lead exposure or the injection (Fig. 8D-E).

**Figure 8.**
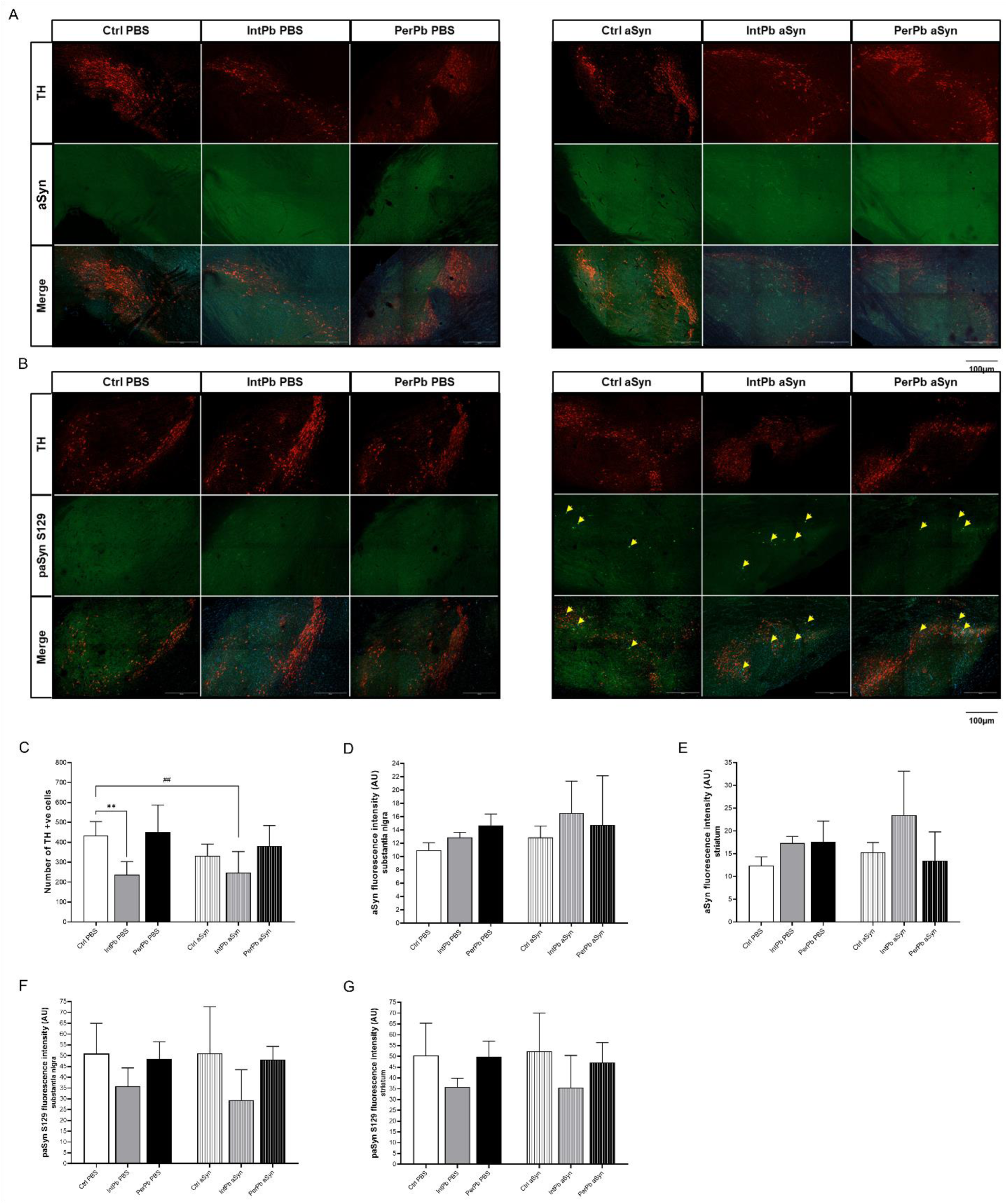
Effect of lead and aSyn PFF injection on dopaminergic neurons (TH-positive cells), aSyn and paSyn S129 levels in the substantia nigra and striatum. A. Representative images of the TH (1:500)-stained dopaminergic neurons and aSyn (1:200) in the substantia nigra. B. Representative images of the TH (1:500)-stained dopaminergic neurons and paSyn S129 (1:200) in the substantia nigra. C. Histogram of the quantification of TH+ve cells. D. Histogram of quantification of aSyn fluorescence intensity in substantia nigra. E. Histogram of quantification of aSyn fluorescence intensity in striatum. F. Histogram of quantification of paSyn S129 fluorescence intensity in substantia nigra G. Histogram of quantification of paSyn S129 fluorescence intensity in striatum. Images were acquired on a confocal point scanning microscope, (Zeiss LSM 900 with Airyscan 2), with a 10x objective. The scale bar is 100 µm for stained images. Values are the mean ± SD for n=4-8 for all groups. The symbols denote statistically significant differences inter groups: Ctrl PBS vs IntPb PBS/PerPb PBS; Ctrl aSyn vs IntPb aSyn/PerPb aSyn - *p < 0.05, **p < 0.01, ***p < 0.001, ****p < 0.0001; Ctrl PBS vs IntPb aSyn/PerPb aSyn; Ctrl aSyn vs IntPb PBS/PerPb PBS - ^#^p < 0.05, ^##^p < 0.01, ^###^p < 0.001, ^####^p < 0.0001; one-way ANOVA, Tuckey’s multiple comparison test.

The morphological analysis of the paSyn-stained images showed that groups injected with aSyn display aSyn inclusion pathology in the *substantia nigra* region (yellow arrows that point to the inclusions), some of which occurs in dopaminergic (TH-positive) neurons (Fig. 8B). We observed no significant differences in paSyn S129 signal in substantia nigra or striatum (Fig. 8F-G).

## Discussion

We are living a PD “pandemic” that is proposed to be associated with the consequences of industrialization, i.e., environmental factors (Dorsey *et al*., 2018). In this context, heavy metals such as lead have been shown to act as toxicants in a variety of biological processes, increasing the risk for several age- and exposure-associated disorders. Several behavioral alterations associated with lead exposure in early life have been reported, including anxiety, locomotor dysfunction and memory impairment, manifesting years after the exposure (Chiodo *et al*., 2004; Toscano & Guilarte, 2005; Sanders *et al*., 2009; Honegger *et al*., 2011; Mason *et al*., 2014; Virgolini & Aschner, 2021; Ortega *et al*., 2021; Shvachiy *et al*., 2023). In the context of neurodegenerative diseases, most previous studies focused on AD, and on the effect of lead on Aβ and tau proteins, on epigenetics/DNA methylation, on protein expression and regulation, and on RNA biology (Bihaqi & Zawia, 2013; Eid *et al*., 2016; Masoud *et al*., 2018). Studies in humans suggested that exposure, even if only for short periods, increases the risk for PD (Gorell *et al*., 2006; Coon *et al*., 2006; Weisskopf *et al*., 2010b; Paul *et al*., 2021). However, the precise molecular mechanisms involved are unclear.

Here, we conducted a broad molecular and physiological assessment of the effect of lead exposure in cell and animal models of PD and related synucleinopathies. Importantly, we established new protocols to mimic different lead exposure profiles - permanent and intermittent low-level lead exposure - in cell cultures. We observed that both lead-exposure paradigms significantly increased aSyn inclusion formation. A similar effect was observed when adding lead to primary neuronal cultures for 16 days (from DIV5 to DIV21) treated with aSyn PFFs (Volpicelli-Daley *et al*., 2014). Nevertheless, we observed a stronger effect in the permanent lead exposure paradigm. Consistent with lead-associated neurotoxicity we observed signs of severe distress with axonal damage, consistent with effects on ion channels, signaling pathways, and mitochondrial function (Garza *et al*., 2006; Zhang *et al*., 2011; Mason *et al*., 2014; Virgolini & Aschner, 2021).

The precise effects of lead on aSyn-associated pathology are still elusive. However, lead causes an upregulation of aSyn and affects apoptosis mechanisms associated with tau pathology (Bihaqi *et al*., 2018). The formation of proteinaceous inclusions in response to lead exposure underscores the possible role of lead exposure in the origin and propagation of aSyn pathology (Murray *et al*., 2001; Iwatsubo, 2007; Goedert *et al*., 2017; Henderson *et al*., 2019b). Consistently, our studies in animals revealed several changes in the presence of lead and/or aSyn injection in the striatum. We analyzed the behavior of at 28 weeks of age, after aSyn injection and with a long-term (28 weeks) lead exposure protocol to analyze the effects of the dual hit - lead and aSyn. We found that lead exposure has a strong effect on readouts of motor behavior, even in the absence of aSyn pathology. However, the strongest effect was observed in episodic long-term memory with a strong impairment observed in the intermittent and permanent lead-exposed groups. Although the effects of lead were observed even in the absence of aSyn pathology, environmental exposure to heavy metals and other toxicants can have an impact on aSyn, promoting more rapid progression of synucleinopathies and other neurodegenerative diseases (Cannon & Greenamyre, 2011; Mason *et al*., 2014; Chin-Chan *et al*., 2015; Nandipati & Litvan, 2016; Villar-Piqué *et al*., 2016).

Interestingly, we observed that all groups of animals treated with lead exhibited strong hypertension, with effects of both systolic and diastolic blood pressure and, consequently, in the mean blood pressure, without changes in the heart rate. Hypertension has been broadly associated with lead toxicity and in some forms of PD (Sharp *et al*., 1987; Gonick *et al*., 1997; Vaziri, 2008; Espay *et al*., 2016; Yoo *et al*., 2021; Grosu *et al*., 2023)Similarly, we also observed tachypnoea and chemoreceptor reflex hypersensitivity in the permanent lead exposure animals, regardless of aSyn injection, suggesting that the effects of lead on autonomic function may have occurred prior to aSyn injection. Likewise, and as previously described, we found that PerPb has a stronger effect on respiratory function and on autonomic control, indicating these animals are in a chronic alert-like state (Shvachiy *et al*., 2018, 2020a, 2022). All cardiorespiratory changes observed are consistent with the blood levels of lead that were observed, with higher levels in the PerPb group which were above the levels considered safe by the World Health Organization (World Health Organization, 2010, 2021; Organization, 2022). Weight changes, which could be a confounding factor in behavioral and physiological alterations, were not present. However, we observed a slight decrease in water intake in the IntPb aSyn group together with a decrease in urine production in the group of animals treated with lead and aSyn, suggesting a slight change in the intake and excretion of lead (Han *et al*., 1999).

In terms of molecular alterations, we observed that lead caused a strong astrocytic activation in the hippocampus. The injection of aSyn did not exacerbate this effect. We found that some cells showed a “ghost-like” morphology, with atrophy of the cells and regression of the processes of the cells, an alteration that has been described before in PD and other diseases like Amyotrophic lateral sclerosis or frontotemporal dementia (Martin *et al*., 2001; Rossi *et al*., 2008; Ramos-Gonzalez *et al*., 2021; Wang *et al*., 2021). These cells have been hypothesized to take up lead prior to neurons, and to store it as a protective mechanism to neurons, similar to what happens with aSyn that is taken up by astrocytes from neuronal cells (Garza *et al*., 2006; Sofroniew & Vinters, 2010; Chen *et al*., 2015; Siracusa *et al*., 2019; Virgolini & Aschner, 2021; Wang *et al*., 2021).

Surprisingly, the levels of total aSyn and of paSyn (widely used as a marker of pathology) were slightly reduced in the hippocampus in animals treated with both lead and aSyn, independently of the type of lead exposure. This effect may be explained by the atrophy and reduced number of the astrocytic cells that we described previously in the hippocampus, compromising the retrieval of pathological forms of aSyn (Pinho *et al*., 2019b; Henderson *et al*., 2019b; He *et al*., 2021). Similarly, we observed a strong degeneration of dopaminergic neurons (TH-positive cells) in the IntPb groups, and a slight decrease in these cells in the presence of aSyn alone in the *substantia nigra*, with a slight increase in the levels of aSyn in this brain region (Spillantini & Goedert, 2000; Murray *et al*., 2001; Goedert *et al*., 2017; Radhakrishnan & Goyal, 2018; Urbina *et al*., 2018; Henderson *et al*., 2019b).

The pattern of paSyn was correlated with the loss of dopaminergic neurons, similar to what happened in the hippocampus. In particular, the IntPb group injected with aSyn showed a decrease in paSyn levels, although we observed visible aSyn inclusions in dopaminergic neurons in the groups injected with aSyn. These observations confirm pathological alterations and suggest aSyn S129 phosphorylation may depend on the pathological insult (Majbour *et al*., 2016; Wang *et al*., 2021). In the striatum, where aSyn PFFs were injected, we observed an increase in the levels of aSyn in the groups injected; we found a trend showing higher levels of aSyn in the IntPb group, but this did not reach statistical significance and will have to be further evaluated in future studies. The distribution of the paSyn signal was similar to that in the *substantia nigra*, with a slight decrease in the IntPb group, possibly due to the loss of the dopaminergic neurons (Paumier *et al*., 2015; Majbour *et al*., 2016; Dehay & Bezard, 2019; Earls *et al*., 2019). It is possible that longer incubation times after aSyn PFF injection might lead to stronger effects, but this will have to be evaluated in future studies as it will involve a different protocol design.

In summary, our results support a mechanistic correlation between environmental lead exposure and the onset and progression of diseases characterized by aSyn pathology accumulation, such as PD. Deciphering the intricate molecular and cellular interactions between lead and aSyn holds substantial implications for informing public health policies. Furthermore, such insights may offer novel perspectives on therapeutic strategies geared towards alleviating the effects of environmental toxicants on neurodegenerative processes, such as those manifesting in synucleinopathies.

## Author Contributions

Conceptualization: L.S., I.R., T.F.O. and V.G. Experimentation: L.S., V.G., Â.A.-L. and F.M. Data analysis: L.S. Manuscript writing: L.S., V.G. and T.F.O. Supervision, V.G., I.R. and T.F.O. Funding: V.G. and T.F.O. All authors have read and agreed with the submitted version of the manuscript.

## Institutional Review Board Statement

The animal experimental protocol was in accordance with the European and national law on animal welfare and was approved by the relevant local ethics committees. The primary cell cultures protocol was approved by the Lower Saxony State Office for Consumer Protection and Food Safety with the approval code 19/3213 and the animal behaviour and manipulation protocol had the approval of the ethics committee of the Academic Medical Center of Lisbon (CAML), Portugal with approval code 411/16.

## Data Availability Statement

The data presented in this study are available on request from the corresponding authors upon reasonable requests.

## Acknowledgements

The authors would like to acknowledge the animal facility from the Faculty of Medicine of Lisbon, Portugal.

